# *Arsenophonus* symbiosis with louse flies: multiple origins, coevolutionary dynamics, and metabolic significance

**DOI:** 10.1101/2023.07.13.548870

**Authors:** Jana Říhová, Shruti Gupta, Alistair C Darby, Eva Nováková, Václav Hypša

**Affiliations:** Department of Parasitology, Faculty of Science, University of South Bohemia, České Budějovice, Czech Republic; Institute of Infection, Veterinary and Ecological Sciences, University of Liverpool, UK; Institute of Parasitology, Biology Centre, ASCR, v.v.i., České Budějovice, Czech Republic

## Abstract

*Arsenophonus* is a widespread insect symbiont with life strategies that vary from parasitism to obligate mutualism. In insects living exclusively on vertebrate blood, mutualistic *Arsenophonus* strains are presumed to provide B vitamins missing in the insect host diet. Hippoboscidae, obligate blood-feeders related to tsetse flies, were previously suggested to have acquired *Arsenophonus* symbionts in several independent events. Based on comparative genomic analyzes of eleven Hippoboscidae-associated strains, nine of them newly assembled, we reveal a wide range of their genomic characteristics and phylogenetic affiliations. Phylogenetic patterns and genomic traits split the strains into two different types. Seven strains display characteristics of obligate mutualists with significantly reduced genomes and long phylogenetic branches. The remaining four genomes cluster on short branches, and their genomes resemble those of free-living bacteria or facultative symbionts. Both phylogenetic positions and genomic traits indicate that evolutionary history of the Hippoboscidae-*Arsenophonus* associations is a mixture of short-term coevolutions with at least four independent origins. The comparative approach to a reconstruction of B vitamin pathways across the available *Arsenophonus* genomes produced two kinds of patterns. On one hand, it indicated the different importance of individual B vitamins in the host-symbiont interaction. While some (riboflavin, pantothenate, folate) seem to be synthesized by all Hippoboscidae-associated obligate symbionts, pathways for others (thiamine, nicotinamide, cobalamin) are mostly missing. On the other hand, the broad comparison produced patterns which can serve as bases for further assessments of the pathways’ completeness and functionality.

**Importance:** Insects that live exclusively on vertebrate blood utilize symbiotic bacteria as a source of essential compounds, e.g. B vitamins. In louse flies, the most frequent symbiont originated in genus *Arsenophonus*, known from a wide range of insects. Here, we analyze genomic traits, phylogenetic origins, and metabolic capacities of eleven *Arsenophonus* strains associated with louse flies. We show that in louse flies *Arsenophonus* established symbiosis in at least four independent events, reaching different stages of symbiogenesis. This allowed for comparative genomic analysis, including convergence of metabolic capacities. The significance of the results is two-fold. First, based on a comparison of independently originated *Arsenophonus* symbioses, it determines the importance of individual B vitamins for the insect host. This expands our theoretical insight into insect-bacteria symbiosis. The second outcome is of methodological significance. We show that the comparative approach reveals artifacts that would be difficult to identify based on a single-genome analysis.

## Introduction

Most insects maintain associations with communities of bacteria, with life strategies ranging from pathogens to obligate mutualists. The generally accepted view assumes that, upon entering the host, the bacteria display characteristics identical or similar to those of their free-living relatives. During the early stage of adaptation, they are distributed in nonspecialized host cells or extracellularly in the gut lumen and rely on both vertical and horizontal transmission to the new host. From the host’s perspective, these bacteria can be pathogens/parasites or facultative symbionts, often of uncertain significance. Some of them may gradually evolve into obligatory mutualists. This process involves several dramatic changes in their biology and genomes. They usually become restricted themselves to specialized host cells and organs (called bacteriocytes and bacteriomes, respectively; (1)), and their transmission to a new host is exclusively vertical, from mother to progeny. However, the most radical change is rapid genomic evolution. Due to the small effective population sizes and relaxed selection, the symbionts undergo a decay process, manifested by shrinking of the genome, due to the loss of unnecessary genes, and shift of the nucleotide composition toward AT bases (2). This process may eventually lead to complete loss of the symbiont and its replacement with a new competent bacterium (3). Such losses and replacements are manifested by incongruent phylogenies of hosts and their symbionts and also by striking genomic differences of the symbionts within a single monophyletic group of the hosts.

Among the bacterial taxa that are best suited for the investigation of symbiogenesis is the gammaproteobacterial genus *Arsenophonus*, which belongs to the most common insect symbionts. Life strategies of its different strains vary from parasites, e.g., reproductive manipulators, to obligate mutualists necessary for the host reproduction and/or development (4). In insects living exclusively on vertebrate blood, *Arsenophonus* manifests both symbiotic lifestyles, i.e., facultative symbiont and obligate mutualist. For example, in triatomine bugs, it is distributed across various tissues in several species, displaying features of a “young” facultative symbiont with uncertain role for the host (5–8). In contrast, *Arsenophonus* strains with the characteristics typical for obligate mutualists were described from sucking lice of the family Pediculidae, and several species of blood-feeding dipterans (Hippoboscidae). The lice-associated *Arsenophonus* possess such dramatically divergent genome that it was initially not recognized as *Arsenophonus* and had been described as separate genus *Riesia* (9, 10). In Hippoboscidae, the situation is much more complex, suggesting a dynamic process of *Arsenophonus* acquisitions and replacements (11). This insect group is one of the blood-feeding families constituting superfamily Hippoboscoidea, together with Glossinidae, Nycteribiidae, and Streblidae (12). Similar to other Hippoboscoidea, they live exclusively on vertebrate blood. While they undergo holometabolous development, their larvae live in the female uterus and are nourished via milk glands (13). Due to this unique system, called adenotrophic viviparity, hippoboscid individuals utilize vertebrate blood as an exclusive dietary source throughout the whole life cycle. As such, they face a shortage of some important dietary components for which they are dependent on their provision by obligate symbiotic bacteria. B-vitamins are the most often suggested candidates for these missing components (14). However, except for two complete *Arsenophonus* genomes from *Melophagus ovinus* and *Lipoptena cervi*, only 16S rRNA or a few other genes are available for the rest of the strains. This hampers both a reliable phylogenetic reconstruction (essential for elucidation of evolutionary history) and evaluation of metabolic capacities in different strains.

The ability to predict metabolic competence is particularly important, as the metabolic role of obligate mutualists in exclusively blood-feeding insects is unclear. The hypothesis that B-vitamins are the main compounds provided by the symbionts to obligate blood-feeders was proposed many decades ago based on experimental works (15–17) and supported later by identification of crucial genes for B-vitamin biosynthesis or even complete operons horizontally transferred to these symbionts (18, 19). However, comparison of studies does not provide a consistent picture of the necessity of different vitamins for blood-feeding hosts. For example, thiamine was shown to be supplemented by *Wigglesworthia* to its host, tsetse fly (20). In contrast, this pathway is disrupted in the *Arsenophonus* strains from the two *Glossina*-related blood-feeders, the hippoboscid species *M. ovinus* and *L. cervi* (21, 22) (and also in the genomes of several louse-associated obligate symbionts; (19, 23–25).

The Hippoboscidae-associated *Arsenophonus*, with multiple independent strains in different stages of evolution (11) provide an ideal system for studying genome changes during the transition towards obligate mutualism and possible evolutionary convergences. To address this process, we reconstruct nine new *Arsenophonus* genomes from eight hippoboscid species representing five genera, *Pseudolynchia*, *Hippobosca*, *Ornithoica*, *Ornithomya* and *Crataerina* (*C. hirundinis* previously classified within genus *Stenepteryx*). Together with other available *Arsenophonus* genomes (including symbionts of hippoboscid species *M. ovinus* and *L. cervi*), we analyze phylogenetic relationships of these bacteria, mainly with the focus on their symbiosis establishment and evolution, i.e., the dynamics of the *Arsenophonus* acquisition across different host species. We use the whole-genome characteristics to evaluate the nature of their symbioses and compare their potential to provide the hosts with essential metabolites.

## Materials and Methods

### Samples preparation

The samples originated from several sources. Some were collected by our lab members in 2012 (*Ornithoica turdi* and *Hippobosca equina*) and 2020 (*Crataerina* spp.). The samples of *Ornithomya* spp. were provided in 2019 by the Department of Zoology (Faculty of Science, University of South Bohemia, Czech Republic). The *Pseudolynchia canariensis* sample was acquired from Dr. Kayce C. Bell (Department of Mammalogy, Natural History Museum of Los Angeles County, Los Angeles, CA, USA). The complete list with collection sites and avian/mammal hosts is provided in the **Supplementary Table 1a, b**. All samples were stored in 96% ethanol at -20 °C. The DNA was extracted using QIAamp DNA Micro Kit (Qiagen) from the abdomen of each individual. DNA quality was assessed by gel electrophoresis, and its concentration measured with a Qubit High sensitivity kit.

### Genomes assembly

All samples were sequenced on the Illumina NovaSeq6000 platform (W. M. Keck Center, University of Illinois at Urbana Champaign, Illinois, USA) generating 2 × 250 paired-end reads. The quality of raw reads was checked using FastQC (26) and the low-quality read ends were trimmed using BBTools (https://jgi.doe.gov/data-and-tools/bbtools). The number of resulting reads for each dataset is provided in **Supplementary Table 1c**. The trimmed reads were assembled using SPAdes 3.10 (27), producing metagenomic assemblies. The bacterial contigs were identified by blasting all genes from *Arsenophonus nasoniae* FIN (NZ_CP038613.1), *Arsenophonus melophagi* (SAMN33924085) and *Arsenophonus lipoptenae* (NZ_CP013920.1) as a query against each metaassembly using custom BLASTn implemented in Geneious Prime v.2020.2.5 program (28). We retrieved five complete *Arsenophonus* genomes from *Ornithomya avicularia* (936,503 bp), *O. biloba*, (874,825 bp), *O. fringillina* (933,061 bp)*, Crataerina hirundinis* (919,248 bp), and *Ornithoica turdi* (599,419 bp), and 4 genome drafts from *Hippobosca equina* (530 contigs), 2 specimens of *Crataerina pallida* (301 and 333 contigs), and *Pseudolynchia canariensis* (355 contigs).

Since the genome of *Arsenophonus* symbiont in *Hippobosca equina* was highly fragmented (530 short contigs), we sequenced this particular sample also using Oxford Nanopore GridIONx5 (W.M. Keck Center, University of Illinois at Urbana-Champaign, Illinois, USA) generating 3,127,655 reads with the length range from 880 bp to 40,881 bp. Their quality was assessed using NanoPack Tools (29) calculating statistics that pointed at overall shorter read length and their lower quality (mean read quality: 14.1, mean read length: 4,506 bp) most probably caused by fragmentation and yield of the used DNA template. The quality trimming was then performed in Filtlong (https://github.com/rrwick/Filtlong). To assemble the *H. equina* Nanopore reads, we employed Canu assembler (30) which yielded 3,894 contigs. The Nanopore filtered reads were mapped onto the assembly using Minimap2 (Li, 2018). The assembly was polished in the following steps. First, we employed consensus calling and polish using Racon (31) and two iterations of Medaka polish (https://github.com/nanoporetech/medaka). Second, to obtain optimal sequence correctness, we mapped Illumina reads onto the assembly using Minimap2 and then we used Racon tool to correct the assembly by consensus generation and polishing. The resulting assembly consisted of 3,595 contigs. The filtering BLAST procedures (as described above) retrieved 16 contigs of the *Arsenophonus* symbiont. All new *Arsenophonus* genomes were annotated using PROKKA v.1.12 (32) and deposited in the GenBank under BioProject accession number PRJNA949118 (**Table 1**). Annotations for the corresponding genomes are deposited in Mendeley Data under the “doi” link https://doi.org/10.17632/8jsd4ds57k.3.

**Table 1.** List of the *Arsenophonus* genomes (ordered by genome size) and their main genomic characteristics.

| Species/strain | Insect host | Genome size (bp) | No. of proteins: annotated/hypothetical | %GC | No. of phages | No. of transposons | No. of mobile elements | No. of ankyrin repeats | No. of invasins | No. of pseudogenes: predicted genes/no ORF predicted | No. of phages by Phaster: intact/incomplete/questionable | Accession (NCBI) |
| --- | --- | --- | --- | --- | --- | --- | --- | --- | --- | --- | --- | --- |
| *A. nasoniae* | *Nasonia vitripennis* | 4987107 | 5555/3198 | 38.7 | 29 | 2 | 0 | 1 | 22 | 1307/374 | 23/14/9 | GCA_004768525.1 |
| *A. Triatominarum* | *Triatoma infestans* | 4721517 | 2118/1481 | 37.8 | 19 | 0 | 0 | 0 | 10 | 4057/1161 | 22/37/25 | GCA_001640365.1 |
| *A. nasoniae* | *Nasonia vitripennis* | 3670548 | 3499/1434 | 37.5 | 17 | 2 | 0 | 1 | 16 | 534/121 | 1/8/2 | GCA_000429565.1 |
| *A. apicola* | *Apis mellifera* | 3639254 | 3143/1176 | 37.8 | 3 | 7 | 0 | 1 | 16 | 687/202 | 8/6/2 | GCA_020268605.1 |
| *A. apicola* | *Apis mellifera* | 3315739 | 3081/1069 | 37.5 | 3 | 6 | 0 | 1 | 15 | 433/110 | 1/7/2 | GCA_903968575.1 |
| *Arsenophonus* sp. | *Entylia carinata* | 3228533 | 3689/1967 | 39.6 | 7 | 0 | 0 | 0 | 5 | 1128/160 | 0/14/1 | GCA_002287155.1 |
| *Arsenophonus* sp. | *Hippobosca equina* | 3226959 | 3080/1049 | 37.5 | 8 | 1 | 0 | 0 | 17 | 589/135 | 8/6/1 | SAMN33923977 |
| *Arsenophonus* sp. | *Crataerina pallida* | 3053565 | 3098/1158 | 38.1 | 6 | 0 | 0 | 0 | 12 | 718/134 | 0/7/0 | SAMN33923976 |
| *Arsenophonus* sp. | *Aleurodicus floccissimus* | 3001875 | 4458/1984 | 37.0 | 2 | 0 | 0 | 0 | 14 | 3436/929 | 8/8/0 | GCA_900343025.1 |
| *Arsenophonus* sp. | *Nilaparvata lugens* | 2953863 | 2658/789 | 37.6 | 2 | 1 | 0 | 0 | 13 | 328/83 | 2/2/1 | GCA_000757905.1 |
| *Arsenophonus* sp. | *Crataerina pallida* | 2844341 | 2805/980 | 37.9 | 6 | 0 | 0 | 0 | 11 | 652/128 | 0/9/0 | SAMN33923975 |
| *Arsenophonus* sp. | *Aphis craccivora* | 2424437 | 3206/1427 | 40.1 | 2 | 0 | 0 | 0 | 3 | 2103/718 | 2/5/2 | GCA_013460135.1 |
| *Arsenophonus* sp. | *Bemisia tabaci* | 2328823 | 2711/869 | 38.3 | 2 | 1 | 0 | 0 | 2 | 1421/339 | 0/1/1 | GCA_902713415.1 |
| *Arsenophonus* sp. | *Bemisia tabaci* | 1860497 | 2411/894 | 36.9 | 1 | 3 | 0 | 0 | 1 | 1596/467 | 0/5/1 | GCA_004118055.1 |
| *Arsenophonus* sp. | *Pseudolynchia canariensis* | 1229579 | 1313/469 | 37.6 | 2 | 1 | 0 | 0 | 5 | 316/54 | 1/1/0 | SAMN33923974 |
| *A. melophagi* | *Melophagus ovinus* | 1155312 | 725/62 | 32.2 | 0 | 0 | 0 | 0 | 0 | 39/21 | 0/0/0 | SAMN33924085 |
| *Arsenophonus* sp. | *Ornithomya avicularia* | 936503 | 660/38 | 24.7 | 0 | 0 | 0 | 0 | 0 | 41/11 | 0/1/0 | SAMN33923973 |
| *Arsenophonus* sp. | *Ornithomya fringillina* | 933061 | 660/37 | 29.4 | 0 | 0 | 0 | 0 | 0 | 34/7 | 0/1/0 | SAMN33923972 |
| *Arsenophonus* sp. | *Crataerina hirundinis* | 919248 | 661/41 | 28.8 | 0 | 0 | 0 | 0 | 0 | 45/14 | 0/1/0 | SAMN33923971 |
| *Arsenophonus* sp. | *Ornithomya biloba* | 874825 | 621/31 | 27.6 | 0 | 0 | 0 | 0 | 0 | 37/13 | 0/2/0 | SAMN33923970 |
| *Arsenophonus* sp. | *Ceratovacuna japonica* | 853149 | 528/29 | 18.5 | 0 | 0 | 0 | 0 | 0 | 42/22 | 0/1/0 | GCA_024349725.1 |
| *A. lipoptenae* | *Lipoptena cervi* | 836724 | 633/33 | 24.9 | 0 | 0 | 0 | 0 | 0 | 27/13 | 0/2/0 | GCA_001534665.1 |
| *Arsenophonus* sp. | *Aleurodicus dispersus* | 663125 | 454/37 | 32.2 | 0 | 0 | 0 | 0 | 0 | 177/114 | 0/1/0 | GCA_900343015.1 |
| *Arsenophonus* sp. | *Ornithoica turdi* | 599419 | 565/23 | 23.6 | 0 | 0 | 0 | 0 | 0 | 27/8 | 0/3/0 | SAMN33923969 |
| *R. pediculicola* | *Pediculus humanus capitis* | 574390 | 480/31 | 28.5 | 0 | 0 | 0 | 0 | 0 | 24/3 | 0/1/0 | GCA_000093065.1 |
| *R. pediculischaeffi* | *Pediculus shaeffi* | 566667 | 496/53 | 31.7 | 0 | 0 | 0 | 0 | 0 | 57/18 | 0/2/0 | GCA_002073895.1 |
| *Riesia* sp. | *Pthirus gorillae* | 528700 | 461/24 | 25.1 | 0 | 0 | 0 | 0 | 0 | 27/6 | 0/1/0 | GCA_002074035.1 |

### Symbiont phylogeny

To determine phylogenetic position of the newly sequenced *Arsenophonus* strains, we downloaded from the NCBI database all available genomes of *Arsenophonus* and four additional gammaproteobacteria as outgroups, *Haemophilus parainfluenzae* (NZ_CP007470.1)*, Sodalis glossinidius* (AP008232), *Proteus mirabilis* (NC_010554.1) and *Providencia stuarti* (NZ_CP095443.1). For this dataset, we built the matrix by concatenating 57 shared single-copy orthologs determined by OrthoFinder (33). The sequences were aligned in MAFFT v.7.450 (34) under E-INS-i settings implemented in Geneious Prime v.2020.2.5 platform (28), and processed by Gblocks (35) using the options for more stringent selection. The resulting matrix of 10,885 amino acids was analyzed by several alternative phylogenetic approaches. Maximum likelihood trees were retrieved by PhyML 3.0 (36) and IQ-TREE-2 (37), Bayesian tree was inferred by PhyloBayes-MPI (38). PhyML analysis was performed under the best fitting model CpREV+R+F selected using Smart Model Selection tool (39) with 100 bootstrap replicates. IQ-TREE-2 analysis was run with 1,000 ultrafast bootstraps under the cpREV+F+I+I+R4 selected by the program according to BIC. Since the data contained taxa with extremely different branch lengths and nucleotide composition, we used PhyloBayes with CAT-GTR model to minimize the artifacts (40). The analysis was run for 30,000 generations. The chain convergence was assessed using the bpcomp and tracecomp commands (checking for the maximum bipartition difference and effective population size of the sample). As an alternative to this standard PhyloBayes analysis, we also analyzed a matrix recoded by Dayhoff6 scheme, which could possibly further decrease the artifacts caused by the composition heterogeneity (41). List of accession numbers for the used genomes and orthologs is provided in **Supplementary Table 1d,e** .The alignment and the phylogenetic trees are deposited in Mendeley Data under the “doi” link https://doi.org/10.17632/8jsd4ds57k.3.

### Host phylogeny and coevolutionary analyses

The phylogeny of hosts was reconstructed for two reasons. First, for the newly assembled samples, it served for the verification of the hosts’ taxonomic determination. Second, it was used to evaluate coevolutionary history and independent acquisitions of the *Arsenophonus* symbionts. For the new samples, we identified mitochondrial genomes by blasting (BLASTn) the complete mitochondrial genome of *Melophagus ovinus* against the whole assemblies. The contigs corresponding to the mitochondrial genomes were then annotated in Mitos server (42). To construct the matrix, we used relevant COI sequences from the new mitochondrial genomes, other COI sequences of hippoboscids available in NCBI, and *Glossina* and *Gasterophilus* as outgroups (**Supplementary Table 1f**). The sequences were aligned by MAFFT v.7.450 under E-INS-i settings implemented in Geneious Prime v.2020.2.5 platform (the alignment did not contain any indels which would require codon-based aligning procedure). The trees were inferred by IQ-TREE-2 (1000 ultrafast bootstrap under the model GTR+F+I+G4 selected by program).

To utilize all information provided by the mitochondrial genomes, we constructed and analyzed another matrix, composed of 12 mitochondrial protein coding genes of the taxa for which complete or nearly complete mitochondrial genomes are available (**Supplementary Table 1g**). The sequences were aligned by MAFFT v.7.450 under E-INS-i settings implemented in Geneious Prime. The matrix was processed in Gblocks using options for more stringent selection. The resulting matrix consisted of 3,205 aa. The concatenated multigene phylogenetic matrix was analyzed using the IQ-TREE-2 (1000 ultrafast bootstrap under mtInv+I+G4 model selected by the program). The alignments and the phylogenetic trees are deposited in Mendeley Data under the “doi” https://doi.org/10.17632/8jsd4ds57k.3.).

### Genome content and metabolic capacity

Since the analyzed genomes were obtained from different sources (our own assemblies and NCBI downloads), we first performed their *de novo* annotation by PROKKA v.1.12 (32) to obtain a standardized input for the downstream analyzes. Based on these annotations, we inferred the main parameters of the genome contents (number of genes, phages, mobile elements, etc., **Table 1**). The number of phages was further verified by PHASTER (43). To estimate number of pseudogenes, we used the Pseudofinder program (44) with default settings. Since some of the symbionts were potentially closely related, we investigated possible synteny of their genomes using two programs, Mauve (45) and Clinker (46).

Metabolic capacities were assessed using the KEGG server (47). K numbers, which assign metabolic functions to the annotated genes, were identified for each genome by the BlastKOALA server (47). Using the KEGG-based structure of metabolic pathways, we mapped these capacities for each genome (**Supplementary Table 2**). Based on this overview, and using KEGG metabolic pathways as a background, we assessed completeness and therefore potential functionality of the pathways for the synthesis of B-vitamins and amino acids.

In order to visualize similarities among the genome contents, we employed MicroEco v0.13.0 (48) package in R environment and performed PCoA (Principal Coordinates Analysis) based on Bray Curtis distances calculated for two datasets, first composed of all orthologs identified by OrthoFinder, and the second composed of genes with assigned K numbers. The graphical outputs were generated utilizing ggplot2 v3.4.0 (49) and further processed in Adobe Illustrator v25.4.1.

## Results and Discussion

The set of *Arsenophonus* symbionts from Hippoboscidae covered a broad range of evolutionary stages, reflected by the genomic characteristics, such as genome size, GC content, presence of mobile genetic elements and branch lengths in the phylogenetic trees. Of the nine new metagenomic assemblies, five produced complete closed *Arsenophonus* genomes (*Ornithomya avicularia*, *O. biloba*, *O. fringillina*, *Crataerina hirundinis*, and *Ornithoica turdi*) and four contained genomic drafts fragmented into contigs (*Hippobosca equina* in 16 contigs, 2 specimens of *Crataerina pallida* in 301 and 333 contigs, and *Pseudolynchia canariensis* in 355). The last genome (from *P. canariensis*) was of insufficient completeness and was therefore excluded from the interpretation of the metabolic analysis, it was however retained in the phylogenetic analyses as it provided a sufficient number of orthologs for building the matrix.

### Arsenophonus *phylogeny*

Phylogenetic analyses revealed several origins of hippoboscid-associated *Arsenophonus*, supporting the results previously obtained by analyzing 16S rRNA genes (11). The trees inferred here by several methods from a multigene matrix agreed on several patterns. The most notable is the split between the small, long-branched genomes and the large, short-branched genomes. Seven of the Hippoboscidae symbionts formed long branches, suggesting that they have experienced a long history of vertical transmission and accelerated molecular evolution, leading to the divergence from the other *Arsenophonus* lineages (**Figure 1**, red taxa; **Supplementary Figure 1).** The remaining four lineages were placed within a cluster of short branches with a poor inner resolution, indicating that these are recent symbioses where the symbionts are highly similar to other *Arsenophonus* lineages from different host groups (blue taxa in **Figure 1**).

**Figure 1.**
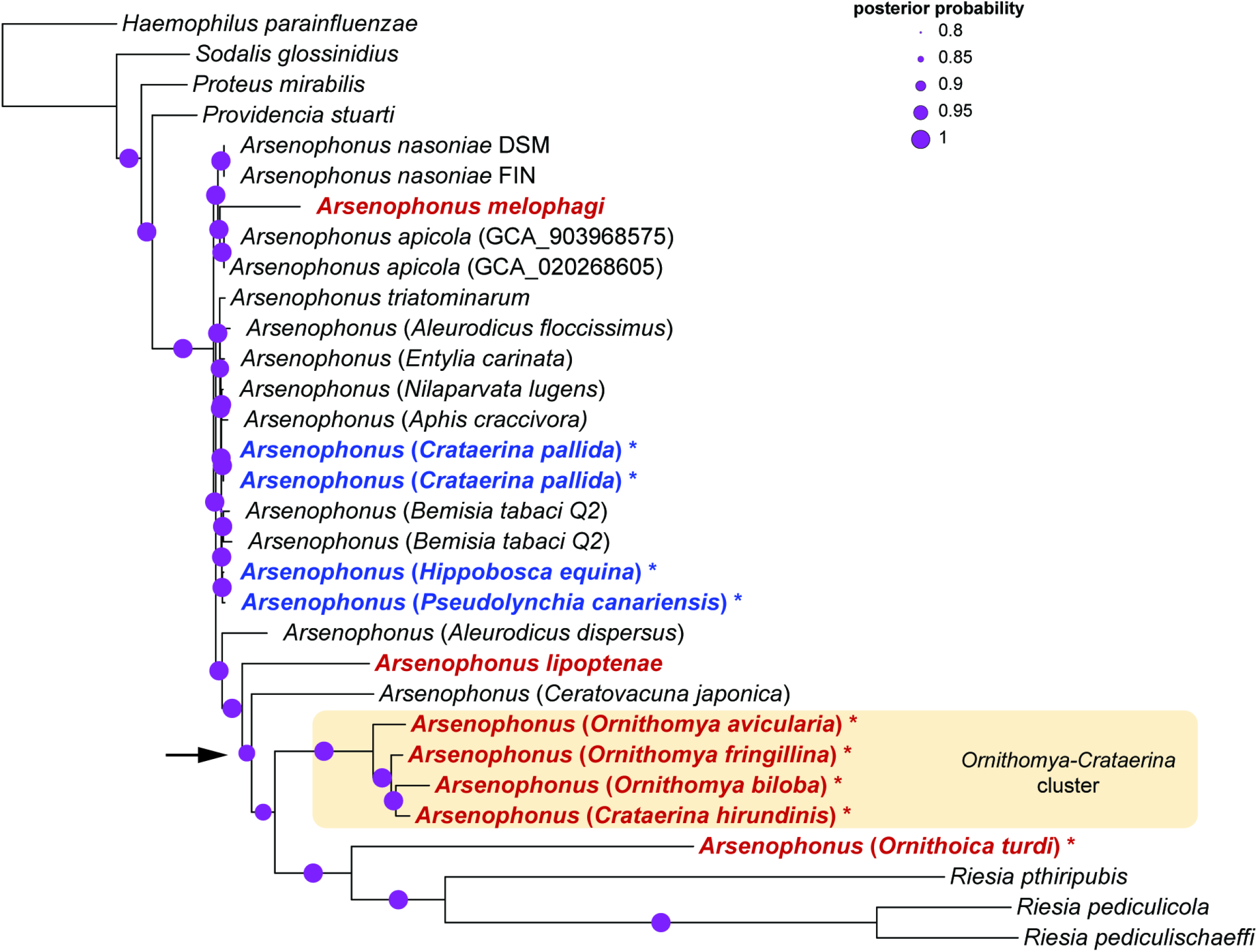
PhyloBayes phylogeny of *Arsenophonus* symbionts derived from concatenated matrix of 57 shared single-copy orthologs (10,885 aa) under CAT-GTR model. The analysis was run for 30,000 generations, maxdiff reached 0.04, and the effective sizes of the parameters were between 531 and 29,910. The hippoboscid-associated taxa considered in the text as “long-branched” and “short-branched” are printed in red and blue, respectively. The asterisks indicate the genomes newly assembled in this study. Arrow indicates the unstable node.

Another stable pattern is the monophyly of four symbionts obtained from three *Ornithomya* species and *Crataerina hirundinis*, which group on a long common branch (*Ornithomya-Crataerina* cluster hereafter). Less clear are the relationships of the remaining three long-branched symbionts (from the genera *Ornithoica*, *Melophagus,* and *Lipoptena*). In the trees, each formed a separate branch with no clear affinity to the other hippoboscid-associated symbionts. However, most of the hippoboscid-associated long-branched *Arsenophonus* symbionts cluster close to the extremely long *Riesia* branch. The topologies of all trees show a clear tendency to cluster according to the branch lengths, which suggests possible role of a long-branch artifact (LBA; 50). Since we applied several approaches which should, in theory, be able to overcome this source of artifact, it is difficult to interpret this pattern. On the one hand, it could result from the LBA despite the algorithms used; on the other it can reflect the real state and tendency of these clusters to establish obligate symbioses.

### Genomes characteristics and evolution

Comparison of the eleven Hippoboscidae-associated *Arsenophonus* strains illustrates typical genomic changes that accompany transition between the recently acquired symbionts (corresponding to the short-branched taxa in the phylogeny, called ‘nascent’ thereafter) and the highly modified types, most likely representing obligate mutualists (the long-branched taxa, called ‘obligate’ thereafter) (**Table 1**). Three of the *Arsenophonus* strains possess relatively large genomes of app. 3 Mbp (one from *Hippobosca equina*, two from *Crataerina pallida*). This size indicates that the strains are likely recently acquired symbionts. Consistent with the genome size are also other genomic characteristics, such as GC content (around 38%) and presence/absence of genes connected to the fluidity of the genomes. For example, all these genomes contain several phage-associated genes and a high number of cell-invasion-associated genes. Although the number of these genes is notably lower than in other ‘nascent’ *Arsenophonus* strains (from *Nasonia*, *Triatoma,* and *Apis*), it sharply contrasts with their complete absence in the seven reduced ‘obligate’ *Arsenophonus* strains. The latter are considerably smaller, but still span a large range from 1.15 Mbp in *Melophagus ovinus* to 0.6 Mbp in *Ornithoica turdi*. In addition, their GC contents range from 32% to 23.6%. Such traits are also mirrored in the phylogenetic pattern. All these genomes form long branches that exceed all other taxa except for the three highly modified genomes of the louse-associated *Arsenophonus* (*Riesia*). Additional indirect supports for their role as obligate, nutritionally essential mutualists comes from the symbiotic system described from the sheep ked, *Melophagus ovinus* (21). In this wingless insect, *Arsenophonus* was shown, using FISH and electron microscopy, to reside in the host bacteriome. Similarly to *Wigglesworthia* in tsetse flies (51), it was also localized in ‘milk glands’, suggesting vertical transmission from mother to progeny via this organ. Among the seven ‘obligate’ Hippoboscidae-associated strains analyzed here, the one from *M. ovinus* possesses the largest genome with the highest proportion of GC. This indicates that the other strains with even smaller genomes and lower GC are also very likely mutualists. Finally, as described below, the obligate mutualistic nature of four of them (*Ornithomya-Crataerina* cluster) is supported by their cophylogeny with the hosts.

The difference in genome characteristics of ‘obligate’ and ‘nascent’ symbionts (**Table 1**), particularly for the repetitive sequences, can likely explain qualitatively different assembly outcomes. For five ‘obligate’ symbionts, we were able to assemble complete closed genomes. Also, of the NCBI-downloaded genomes, the smaller one, from *Lipoptena*, is available in a single complete contig. In contrast, for the ‘nascent’ symbionts we could only recover fragmented assemblies, despite the fact that for *Hippobosca equina* sample, we also generated long Nanopore reads (see Materials and Methods). The *Arsenophonus* strain from *P. canariensis* possessed a seemingly mixed set of characteristics. In the phylogeny, it clustered on a short branch, among other ‘nascen’ strains. It contained phage-related sequences, and its GC content (37.6 %) was also close to the other ‘nascent’ symbionts. On the other hand, the size (1.23 Mbp) resembled that of the ‘obligate’ symbionts (**Table 1**). However, we presume that the genome size is the result of an incomplete assembly rather than a reductive evolutionary process. This view is the simplest (most parsimonious) explanation of the observed conflict and is further supported by the notable inconsistency of the metabolic reconstruction when compared to the other analyzed genomes (**Supplementary Table 2**).

The split between ‘obligate’ and ‘nascent’ strains is well reflected in the PCoA analyses of the two genome-content characteristics (shared orthologs and shared genes with assigned K numbers). Both analyses showed clear separation between the two symbiont types along the main axis, which explains a very high portion of the variability (75.5% and 78.7%). The gap between these two forms is interesting and fits the general view on the evolutionary tempo during symbiogenesis. It assumes that once established in the host, the bacteria experience a period of rapid genomic changes (mainly degeneration), after which the tempo of the changes slows down, potentially reaching almost complete stasis (52–55). Therefore, it may be more difficult to capture a genome in the fast-evolution phase, rather than in the stages close to the free-living ancestor or the obligate symbionts (although an artifact due to possibly unrepresentative number of 26 strains cannot be excluded). Apart from this separation, the analyzes yielded additional interesting patterns. The seven ‘obligate’ strains associated with hippoboscids group within a common cluster (orange ellipses in **Figure 2**), although they are scattered in a non-monophyletic manner across the phylogenetic tree (**Figure 1**). In contrast, the three *Riesia* strains (pink ellipses in **Figure 2**), while phylogenetically closely related to the *Ornithomya-Crataerina* cluster, form a distinct and relatively distant group (on the second axis) from the hippoboscid strains. This may imply a convergent adaptation of unrelated strains to particular host groups (hippoboscids vs. lice). On the other hand, this conflict between phylogeny and PCoA clustering could also, in theory, be due to artificial phylogeny, distorted by LBA.

**Figure 2.**
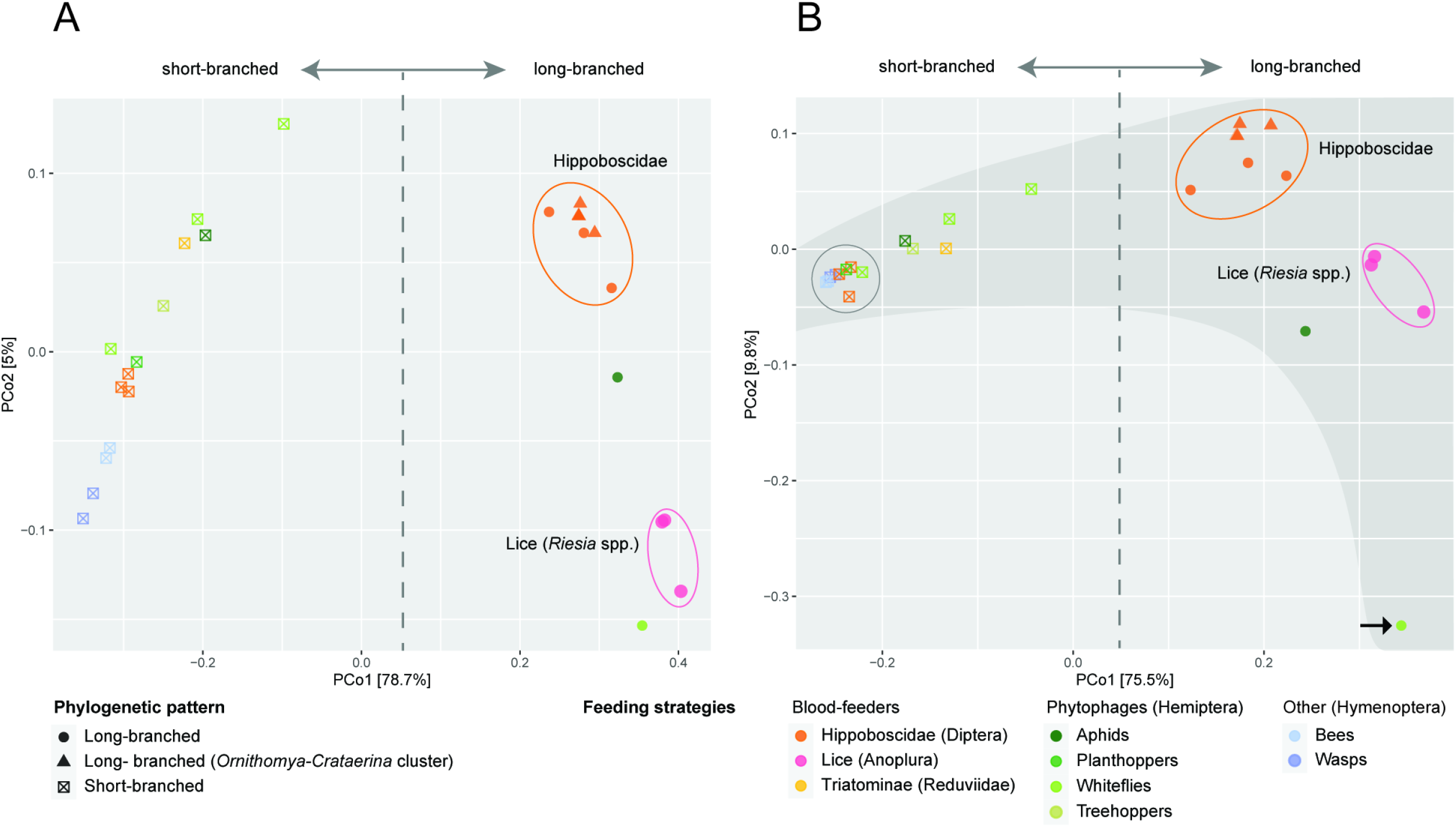
PCoA analyses of genome contents calculated using Bray Curtis distances among all protein coding genes (A) and genes with assigned K numbers (B) found across 26 analyzed *Arsenophonus* genomes. Ellipses (not statistical) show long-branched (obligate) strains associated with Hippoboscidae (orange) and lice (pink). The arrow points to the outlier (see text). The grey-shaded area is explained in the text.

The clustering patterns derived from the two different criteria (shared orthologs and shared K numbers) differed significantly on the second axis. In ortholog-based analysis (**Figure 2A**), the ‘nascent’ and ‘obligate’ strains were broadly distributed, indicating a considerable degree of variability. In the analysis based on K numbers (**Figure 2B**), the ‘nascent’ strains aggregated in a narrow strand, while the ‘obligate’ strains were broadly distributed (indicated by the dark gray background). This pattern shows that ‘nascen’ strains share significantly higher number of the metabolically important genes with assigned functions (i.e. K numbers; **Figure 2A**) than all identified orthologs (including ‘non-necessary’ genes). The difference between the ‘nascent’ and ‘obligate’ strains on the second axis in the analysis based on K numbers fits well with the general view of the evolution of symbionts. While ‘nascent’ symbionts with large genomes possess similar metabolic capacities, ‘obligate’ symbionts exhibit much greater variability, likely due to their adaptation to different conditions and metabolic demands. Since in **Figure 2B** the broad distribution of ‘obligate’ symbionts could be caused by the presence of an outlier (*Aleurodicus dispersus*; indicated by an arrow), we performed another analysis without this genome. Its result (supplementary **Figure 2**) shows a pattern similar to that in **Figure 2B**.

### Coevolution with the host

To estimate the coevolutionary pattern in the *Arsenophonus*-Hippoboscidae association, we compared the *Arsenophonus* tree with the phylogeny derived from the whole mitochondrial genomes available for the hosts (**Figure 3**). The relationship between the Hippoboscidae and *Arsenophonus* phylogenies was consistent with the general genomic characteristics of the symbionts. The only part of the *Arsenophonus* tree that shows a clear sign of cospeciation is the monophyletic lineage composed of long branched strains of the *Ornithomya-Crataerina* cluster. This hippoboscid phylogenetic lineage also provides an example of the complex processes involving both symbionts’ cospeciation and replacement. At the host site the branch is composed of five taxa (two species of *Crataerina* and three species of *Ornithomya*). However, only *Ornithomya* and *C. hirundinis* carry these *Arsenophonus* ‘obligate’ strains. On the contrary, two samples of *C. pallida* harbor *Arsenophonus* with large genome and characteristics typical for ‘nascent’ symbionts. In the *Arsenophonus* phylogeny, this strain clusters among the other ‘nascent’ symbionts of several insect hosts. This arrangement suggests the establishment of the symbiosis prior to the *Ornithomya*-*Crataerina* diversification, followed by cospeciation of the symbiont with the *Ornithomya*-*Crataerina* cluster, and a replacement of this strain with a new phylogenetically distinct *Arsenophonus* in *C. pallida* (**Figure 3**). The correct phylogenetic placement of our samples in the hippoboscid phylogeny/taxonomy was further confirmed by the additional analysis based on the COI gene available for a broader taxonomic sample (**Supplementary Figure 3**).

**Figure 3.**
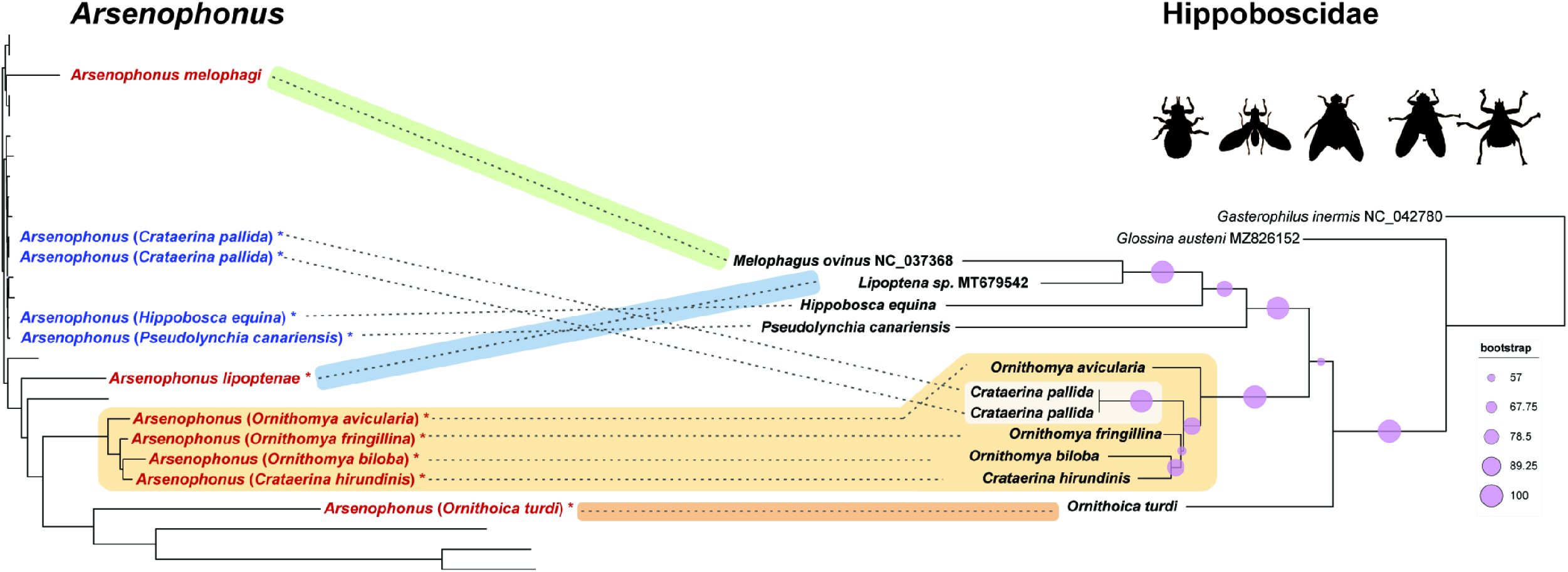
Coevolutionary analysis of the *Arsenophonus* strains and their hippoboscid hosts. **Left**: *Arsenophonus* tree simplified from the Figure 1 (only Hippoboscidae-associated strains are shown). **Right** Hippoboscidae tree derived from matrix of 12 concatenated mitochondrial genes (3,205 aa) in IQ-TREE-2 under the mtInv+I+G4 model. The four presumably independent establishments of the symbiosis are indicated by different colors highlighting the host-symbiont connections. The connections without highlights indicate an uncertain evolutionary interpretation.

Less clear is the evolutionary history of the other ‘obligate’ strains, i.e. the symbionts of *Ornithoica*, *Lipoptena*, and *Melophagus*. In the phylogenetic analyses, the ‘obligate’ strains associated with hippoboscids never formed a monophyletic branch. This arrangement could in theory be affected by long branch attraction, since in some analyses the *Ornithoica*, *Lipoptena*, and *Ornithomya*-*Crataerina* strains did form a paraphyletic group with respect to extremely long-branched *Riesia*. However, based on several phylogenetic and genomic patterns described below, we assume that the four mentioned lineages (*Melophagus*, *Lipoptena*, *Ornithoica*, and *Ornithomya*-*Crataerina*) originated independently within the genus *Arsenophonus*. An independent origin can be most robustly supported for the *Melophagus* symbiont. While located on a long branch, this strain invariantly falls into a monophyletic group with four ‘nascent’ strains from *Nasonia vitripennis* and *Apis mellifera*, at a position phylogenetically distant from the cluster formed by several long branches. The position of the remaining three lineages (*Lipoptena*, *Ornithoica*, *Ornithomya*-*Crataerina*) varied with the phylogenetic analyses, but they never formed a monophyletic clade. The phylogenetic arrangement of these three lineages was the least stable and supported part of the whole phylogeny (**Supplementary Figure 1**). Their independent origins are supported by comparisons of their genome arrangements. It is well known that bacterial genomes are fluid, and due to the presence of various categories of genes (such as phage-associated genes, mobile elements, etc.) can rapidly reorganize their gene content and lose genomic synteny (56). This is true for free-living bacteria and symbionts in the early stage of evolution. In contrast, once they get rid of the fluidity-associated genes, the symbionts lose the rearrangement capacity and retain a high degree of synteny (52, 57). In our set, all ‘obligate’ strains (including those from *Ceratovacuna japonica* and *Aleurodicus dispersus*) lack phage-associated genes as well as transposons, in contrast to the ‘nascent’ strains (**Table 1**). However, only the four strains of the *Ornithomya*-*Crataerina* cluster show a high degree of genome similarity and synteny (**Figure 4**). Despite this, their genomes are not identical, but differ in gene content. This provides evidence that the genomes reached the point when they are not capable of gene rearrangements and they only continue to lose genes. In particular, the *O. biloba* symbiont possesses a genome considerably smaller than the other three strains. Among the genes which this smallest genome lacks is for example the complete pathway for heme synthesis, present in the other three genomes (**Figure 4B**). The synteny within the *Ornithomya*-*Crataerina* cluster contrasts strikingly with the other three lineages of the ‘obligate’ symbionts, for which no sign of synteny could be found. The lack of synteny indicates two possible scenarios. First, the four linages represent four independent origins of *Arsenophonus* symbiosis. Second, they share common, possibly facultative, ancestor, and their diversification into the four non-syntenic lineages had taken place before they lost the fluidity-associated genes. Taking together all of the phylogenetic and genomic evidence, we consider the first scenario much more likely.

**Figure 4.**
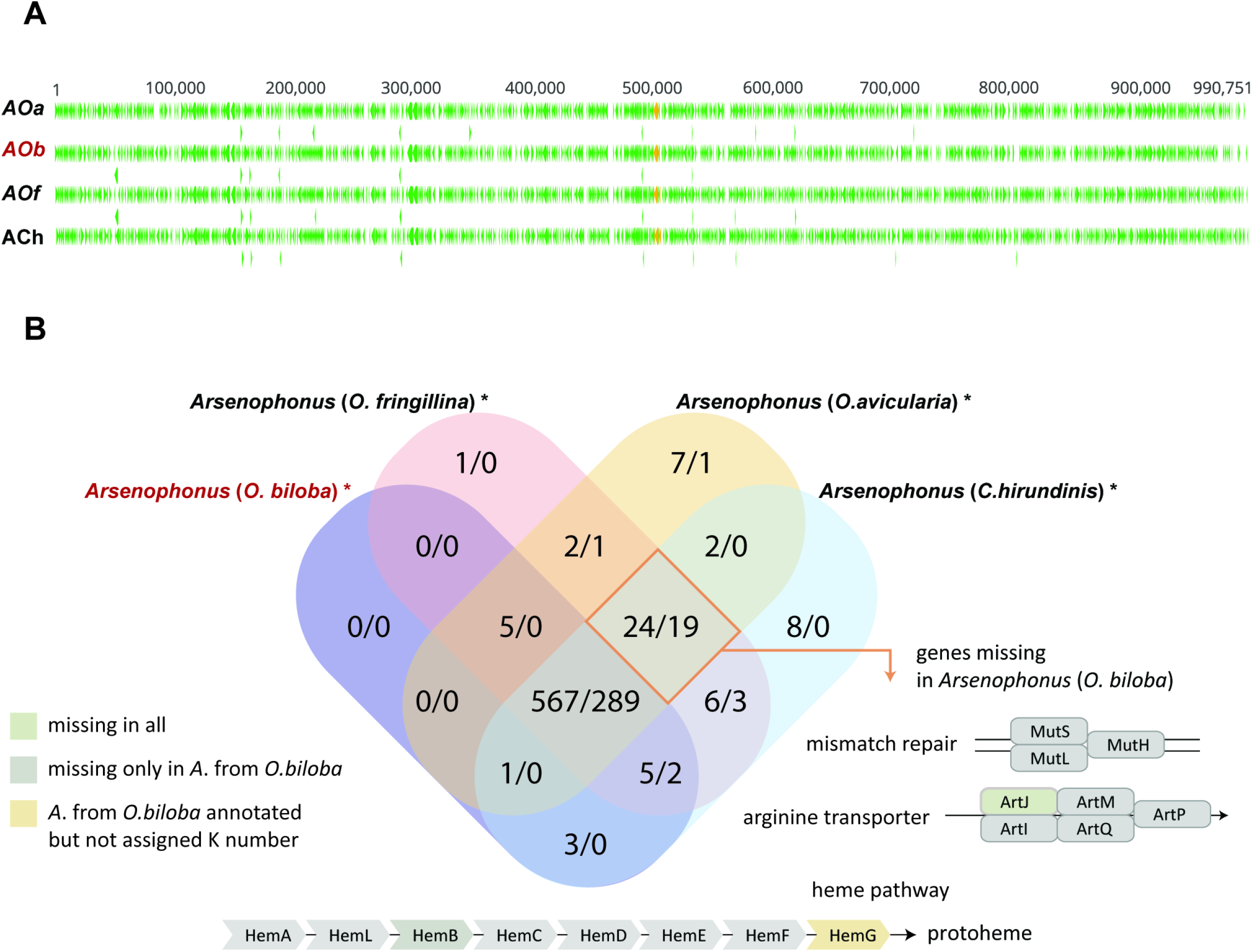
Genomic comparison of the four strains of the *Ornithomya-Crataerina* cluster. **A:** Synteny of the genomes (alignment built by MAFFT algorithm). Strain abbreviations are composed of *A* (*Arsenophonus*) and the host, *Oa* (*O. avicularia*), *Ob* (*O.biloba*), *Of* (*O.fringillina*), and *Ch* (*C. hirundinis*). *Aob* is printed in red to indicate the shortest genome. Green triangles represent protein-coding genes (the genes shown in the second row of each genome overlap with the first-row genes), orange triangles stand for rRNA genes. **B:** Venn diagram of shared genes among the four strains of the *Ornithomya*-*Crataerina* cluster. *Arsenophonus* (*O. biloba*) is printed in red to indicate the shortest genome. Its difference (missing genes) compared to the other strains is indicated by the orange frame. Structures of the heme pathway, mismatch repair, and arginine transporter are adopted from the KEGG database.

### Metabolic capacity of the Arsenophonus symbionts

The overview presented in **Figure 5** (details in **Supplementary Table 2**) shows that the pathways of the eight B-vitamins differ in their completeness across the *Arsenophonus* genomes. Only two of them, riboflavin and lipoic acid, are complete and therefore likely functional in all genomes. Other widely present pathways, but absent or likely incomplete in several genomes, are those for folate, biotin, and pyridoxal synthesis. The most interesting patterns were found for the thiamine and, particularly, pantothenate pathways. The thiamine pathway is clearly not functional in all ‘obligate’ strains (i.e., the obligate symbionts), except for the strain from *Ornithoica turdi*. In this small genome, the *thiI* annotation was assigned to a short-sequence, but it was not recognized as a functional gene by BlastKOALA. Compared to other strains, the annotated gene was extremely shortened (288 bp vs. 1,449-1,485 bp). While the absence of a single gene potentially makes a whole pathway non-functional, this is likely not the case in the strain *Ornithoica turdi* thiamine pathway. First, the presence of the additional nine genes in such a reduced genome (compared with their absence in the other ‘obligate’) suggests a selection in favor of preserving the pathway. Second, several sources of evidence suggest that complete *thiI* may not be necessary for the functionality of the thiamine pathway. Martinez-Gomez et al. (58) showed experimentally that only part of the *thiI* gene, the rhodanese domain, is necessary for the thiamine biosynthesis in *Salmonella*. Using the conserved domain database (59), we revealed that the retained fragment of the *thiI* corresponds to the rhodanese domain (**Figure 5**). A similar situation, that is, preservation of the rhodanese domain instead of the complete *thiI* gene, was reported at least from two other obligate symbionts, *Wigglesworthia glossinidia* from tsetse fly *Glossina morsitans* (60), and *Candidatus* Pantoea from brown marmorated stink bug (61). However, in other cases the pathway exists without a rhodanese domain containing ORF. In comparison of six *Rhodococcus* species and *Micrococcus luteus,* all genomes lacked the *thiI* rhodanese domain while retaining the rest of the thiamine pathway (62). This contrast shows that our knowledge on the synthesis of thiamine by insect symbionts (or generally bacteria) is currently incomplete. Interestingly, consistent with the preserved thiamine pathway, the strain from *Ornithoica* genome is also the only one among ‘obligate’ strains which does not encode for a thiamine transporter. The only other genomes with a missing or incomplete thiamine transporter are *Arsenophonus* strains from *Ceratovacuna japonica* and *Aleurodicus dispersus* which lack all three genes, and *Aleurodicus floccissimus* with one missing gene (**Figure 5, Supplementary Table 2**).

**Figure 5.**
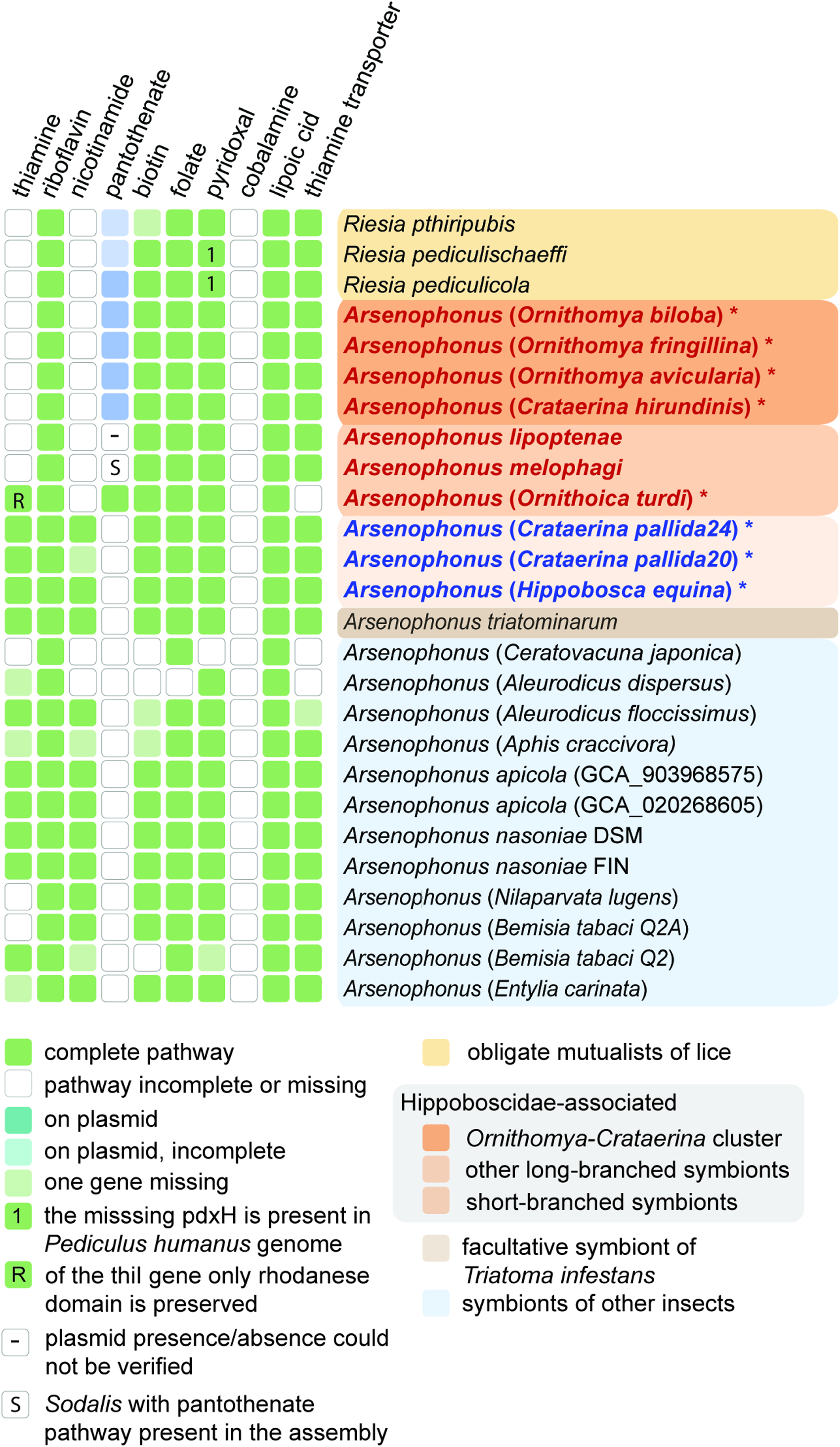
An overview of B-vitamin biosynthetic capacities in the analyzed genomes. The completeness of the pathways was derived by assigning K numbers in BlastKOALA and comparing the results with the structure of the pathways in KEGG (see Methods). *Arsenophonus* strains associated with hipoboscids are printed in bold (red = long-branched/obligate, blue = short-branched/nascent), asterisks = genomes newly assembled in this study.

The distribution and origin of the three genes required for pantothenate biosynthesis (*panB*, *panC*, *panE*) proved to be the most complex and biologically interesting. Most of the analyzed *Arsenophonus* genomes do not possess these genes. The absence of pantothenate-synthesis pathway genes in several of these *Arsenophonus* genomes was previously reported from both phytophagous and hematophagous hosts (21, 63, 64). This absence suggests that pantothenate biosynthesis may have been missing at the origin of the entire *Arsenophonus* clade (*Arsenophonus* synapomorphy). In contrast to the general absence of the pantothenate pathway in *Arsenophonus* bacteria, the four closely related hippoboscid obligate symbionts from the *Ornithomya*-*Crataerina* cluster and the strain from *Ornithoica turdi* possess this metabolic capacity. However, the origin and location of these genes differ among the symbionts. In *Riesia*, the genes have been known to reside on plasmids (25, 65). In this study, we found similar arrangements in four ‘oblgate’ *Arsenophonus* strains from the monophyletic *Ornithomya*-*Crataerina* cluster (**Figure 5**). All of these symbionts possess plasmids with sizes from app. 12kbp to 21kbp, which carry the three pantothenate genes. A different picture was found in the ‘obligate’ symbiont from *Ornithoica turdi*. Here, all three pantothenate genes are located directly on the bacterial chromosome.

Based on this overview, the vitamin pathways in the ‘obligate’ *Arsenophonus* strains can be characterized with four categories in respect to their importance to the host.

#### Non-essential pathways

Two of the vitamins, nicotinate and thiamine, are obviously not crucial, and most of the ‘obligate’ symbionts here lack their synthesis capacity. The retention of thiamine transporter indicates that they can scavenge this vitamin either from the host or from other bacteria, as suggested in several previous studies (18, 21, 66).

#### Universal pathways

Riboflavin and lipoic acid are synthesized by all included strains. This omnipresence makes it difficult to decide if these vitamins are required by hosts or their synthesis is essential for *Arsenophonus* cells.

#### Majority pathways

The biotin, folate, and pyridoxal pathways are present in most of the strains. Folate is only missing in the considerably degenerated strain from *Aleurodicus dispersus* but present in all ‘obligate’ symbionts. Similarly, pyridoxal is only missing the degenerated strain from *Ceratovacuna japonica*. It is also seemingly incomplete in *R. pediculicola and R. pediculischaeffi*. However, the missing gene responsible for the terminal conversion of pyridoxine phosphate to pyridoxal phosphate is present in the *Pediculus* host genome (KEGG code Phum_PHUM170130). Biotin pathways are incomplete in several ‘nascent’ symbionts and possibly in *R. phthiripubis*.

#### Essential pathway

Pantothenate is the only vitamin whose importance for the *Arsenophonus*-Hippoboscidae symbiosis is strongly supported by its complex distribution pattern. It is absent in all ‘nascent’ symbionts but present in all of the Hippoboscidae-associated ‘obligate’ symbionts for which metagenomic data are available (in several cases coded on plasmids). In *A. lipoptenae* and *A. melophagi* the pantothenate pathway is not encoded by the available genomes. In our previous work (22, 67), we did not find plasmids for these two *Arsenophonus* strains. However, we reported that apart from this obligate symbiont, the host *Melophagus ovinus* harbors two additional symbionts, *Sodalis* and *Bartonella*, both capable of pantothenate synthesis (21). Unfortunately, plasmid absence could not be verified in this study due to the unavailability of the raw data. An interesting situation was encountered in the three *Riesia* strains. All three pantothenate genes are annotated on the *R. pediculicola* USDA plasmid. On the basis of the PROKKA annotation and BlastKOALA assignments, the plasmids of the other two strains carry only *panB* and *panC*, while they lack *panE*. The DNA sequence corresponding to this locus is present in both plasmids, but it is highly modified and therefore is not recognized as a functional *panE* homolog (**Figure 6**). Compared to B-vitamins, the pathways for amino acid synthesis are greatly deteriorated in all ‘obligate’ symbionts, while mostly preserved in the ‘nasent’ strains (**Supplementary Table 2**).

**Figure 6.**
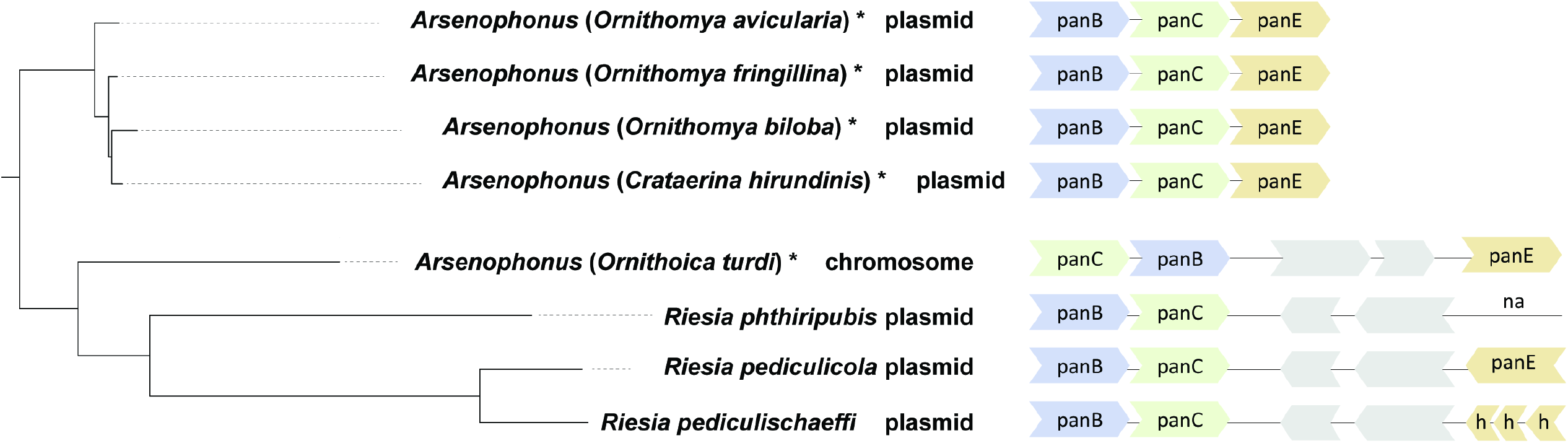
Arrangement of pantothenate genes. Grey: other genes, not involved in the pantothenate pathway; na: not annotated (not recognized as a gene) by PROKKA; h: hypothetical protein.

### Conclusion

This study utilized Hippoboscidae-associated strains of *Arsenophonus* as a convenient system for studying adaptation of bacteria to mutualistic relationship with obligate blood-feeders. In direct relation to the model used, the study produced several concrete findings. For example, the evolutionary history of the Hippoboscidae-*Arsenophonus* association is a mixture of two processes. On the one hand these associations resulted from multiple *de novo* establishments of symbiosis; on the other hand in some clusters they experience a period of coevolution with the hosts. The strains which adapted to obligate mutualism, while originating from different phylogenetic lineages, display notable metabolic convergence, particularly in B-vitamin pathways.

At the general level, in relation to reconstructions of metabolic roles in the obligate blood-feeders, it revealed several weaknesses/problems and also the advantages of such a comparative approach. The first weakness derives from an uncertainty when determining and annotating the genes. When comparing our annotations with other published studies, we occasionally found differences, usually related to highly modified genes (e.g., panE in two species of Riesia; 18, 25). We have observed similar discrepancies even within our study between the functional annotations obtained by different programs (e.g., PROKKA vs. BlastKOALA vs. Pseudofinder). This problem is particularly significant in strains with strongly degenerated genomes, where the genes could be shortened, dramatically changed in nucleotide composition, or pseudogenized. Another weakness stems from our incomplete knowledge of the structure of the pathways. It has been shown in various studies that the functional flexibility of some pathways may be higher than expected. For example, some missing enzymes may not hamper the pathway functionality (68–70), others may be replaced with different enzymes (71, 72) sometimes with so called promiscuous enzymes (73). In contrast, the advantage of a comparative study (apart from providing background for evolutionary interpretation) lies in the indication of possible annotation artifacts, which would be difficult to identify based on single-genome analysis. In our data, we have observed complete pathways lacking the same single enzyme in several taxa (Supplementary Table 2, numbers highlighted by orange); for example tenI (row 6 in Supplementary Table 2), yigB (row 14), epd (row 19), nudB (row 48). For all these genes, it has been previously demonstrated that they are either not essential for the pathway functionality, or can be replaced with different enzymes (69, 72, 74, 75). This strongly suggests that the pathway is conserved by selection with the exception of the missing gene. Such indicated genes and pathways can then be analyzed in more detail, which will lead to better understanding of the degenerative processes during the adaptation to obligate symbiosis.

## Data availability

The genome assemblies of *Arsenophonus* symbionts are available from GenBank under BioProject number PRJNA949118 (individual accessions are provided in Table 1). The mitochondrial genomes of corresponding Hippobocidae are available from GenBank under accession number xxxxxx. The alignments (in fasta format), phylogenetic trees (in newick format) and annotations of *Arsenophonus* genomes (in gbk format) are deposited in Mendeley Data under the “doi” link https://doi.org/10.17632/8jsd4ds57k.3.

## Acknowledgements

Access to computing and storage facilities owned by parties and projects contributing to the National Grid Infrastructure MetaCentrum provided under the programme “Projects of Large Research, Development, and Innovations Infrastructures” (CESNET LM2015042), is greatly appreciated. We thank Greg DD Hurst for his valuable comments and edits to the manuscript.

## Supplementary Material

**Supplementary Figure 1.** *Arsenophonus* trees derived from the 10,885 amino acid matrix by Bayesian inference and ML. A: Phylobayes (30,000 generations), matrix recoded according to Dayhoff6 scheme. B: PHYML under CpREV+R+F model with 100 bootstrap replications. C: IQ-TREE under model cpREV+F+I+I+R4 selected by the program (1,000 samples for ultrafast bootstrap). D: Q-TREE under mixture model JTT+C60+F+R4 (1,000 samples for ultrafast bootstrap). Sequences generated in this study are printed in bold.

**Supplementary Figure 2.** PCoA analyses of genome contents calculated using Bray Curtis distances among all protein coding genes (A) and genes with assigned K numbers (B) found across 26 analyzed *Arsenophonus* genomes. Ellipses (not statistical) show obligate (long-branched) strains associated with Hippoboscidae (orange) and lice (pink). The grey-shaded area is explained in the text.

**Supplementary Figure 3.** Host phylogeny derived from mitochondrial COI gene matrix in IQ-TREE-2 under the GTR+F+I+G4 model with 1000 ultrafast bootstrap. Sequences generated in this study are printed in bold.

**Supplementary Table 1a.** List of accession numbers for the complete genomes and genome drafts of *Arsenophonus* symbionts retrieved in this study (including the samples metadata).

**Supplementary Table 1b.** List of accession numbers for the mitochondrial genomes of hippoboscids retrieved in this study (including the samples metadata).

**Supplementary Table 1c.** Number of trimmed reads for each of the sequenced sample. **Supplementary Table 1d.** List of accession numbers and abbreviations for genomes used in the *Arsenophonus* matrix.

**Supplementary Table 1e.** List of single-copy ortholog genes used in the *Arsenophonus* matrix. The order of genes shown in this table corresponds to the order of gene alignments in the concatenated phylogenetic matrix.

**Supplementary Table 1f.** List of accession numbers and abbreviations for the mitochondrial genomes used in the host matrix.

**Supplementary Table 1g.** List of single-copy ortholog genes used in the concatenated host matrix. The order of genes shown in this table corresponds to the order of gene alignments in the concatenated phylogenetic matrix.

**Supplementary Table 2.** Metabolic overview

